# Fine-scale spatial genetic structure across the species range reflects recent colonization of high elevation habitats in silver fir (*Abies alba* Mill.)

**DOI:** 10.1101/2021.05.02.442307

**Authors:** Enikő I. Major, Mária Höhn, Camilla Avanzi, Bruno Fady, Katrin Heer, Lars Opgenoorth, Andrea Piotti, Flaviu Popescu, Dragos Postolache, Giovanni G. Vendramin, Katalin Csilléry

## Abstract

Variation in genetic diversity across species ranges has long been recognized as highly informative for assessing populations’ resilience and adaptive potential. The spatial distribution of genetic diversity, referred to as fine-scale spatial genetic structure (FSGS), also carries information about recent demographic changes, yet it has rarely been connected to range scale processes. We studied eight silver fir (*Abies alba* Mill.) population pairs (sites), growing at high and low elevations, representative of the main genetic lineages of the species. A total of 1368 adult trees and 540 seedlings were genotyped using 137 and 116 single nucleotide polymorphisms (SNPs), respectively. Sites revealed a clear east-west isolation-by-distance pattern consistent with the post-glacial colonization history of the species. Genetic differentiation among sites (*F_CT_*=0.148) was an order of magnitude greater than between elevations within sites (*F_SC_*=0.031), nevertheless high elevation populations consistently exhibited a stronger FSGS. Structural equation modeling revealed that elevation and, to a lesser extent, post-glacial colonization history, but not climatic and habitat variables, were the best predictors of FGSG across populations. These results may suggest that high elevation habitats have been colonized more recently across the species range. Additionally, paternity analysis revealed a high reproductive skew among adults and a stronger FSGS in seedlings than in adults, suggesting that FSGS may conserve the signature of demographic changes for several generations. Our results emphasize that spatial patterns of genetic diversity within populations provide complementary information about demographic history and could be used for defining conservation priorities.

## Introduction

Dispersal capacity of organisms plays a fundamental role in the establishment, persistence and range dynamics of species, especially during environmental changes (Travis et al., 2013, Saastamoinen et al., 2018). Most terrestrial organisms have limited dispersal, which leads to a non-random spatial distribution of genes and genotypes. The short-term consequence of this process is the clustering of related individuals, which is manifested as a decay-of-kinship with distance, and is commonly referred to as fine-scale spatial genetic structure (FSGS) (Wright, 1943; Epperson, 1995; Sokal & Wartenberg, 1983; Rousset, 2000). The long-term consequence of limited dispersal is that populations become subdivided and distant populations become genetically more differentiated than nearby populations; a phenomenon referred to as isolation-by-distance (IBD) in subdivided populations (Wright, 1943; Malécot, 1948; Kimura & Weiss, 1964). Thus, FSGS captures principally the effect of limited dispersal with respect to the past few generations, while, in contrast, IBD reflects deeper demographic time scales. This distinction is important because, ignoring mutations, a drift-dispersal equilibrium can be reached within a few generations on a spatial scale up to one order of magnitude larger than the average parent-offspring distance, while many more generations can be necessary on larger spatial scales (Slatkin, 1993; Vekemans & Hardy, 2004; Bradburd & Ralph, 2019).

IBD is an ubiquitous property of most biological systems, yet this pattern is driven by many ecological and evolutionary factors, and not only variation in dispersal (Slatkin, 1985; Rousset, 2000; Legendre & Fortin, 2010; Duforet-Frebourg & Blum, 2014). Most species evolutionary histories in the northern hemisphere are dominated by recent colonization dynamics from glacial refugia since the last glacial maximum (LGM) (Petit et al., 2008; Excoffier et al., 2009). In these cases, IBD has little to do with the presence or absence of a drift-migration equilibrium (Meirmans, 2012; Aguillon et al., 2017) while patterns of FSGS across the species range could provide complementary information by elucidating the recent demographic history. Indeed, recently expanded populations are expected to more likely show spatially aggregated patterns of related individuals with respect to demographically old and dense populations (Gapare & Aitken, 2005; De-Lucas et al., 2009; Pandey & Rajora, 2012; Torroba-Balmori et al., 2017). However, patterns of FSGS may also differ from one population to another across the species range for reasons other than the demographic age of populations. Differences may reflect natural or human disturbances, such as management or habitat degradation (e.g. Dreier et al., 2014; Bessega et al., 2016; Piotti et al., 2013), fire (Budde et al., 2017), presence or absence of dispersers (Barluenga et al., 2011; Rico & Wagner, 2016; Gelmi Candusso et al., 2017; Kling & Ackerly, 2021), or adaptation to the local environment (e.g. Troupin et al., 2006; Audigeos et al., 2013; Mosca et al., 2018; Gauzere et al., 2020).

Currently, there is a clear dichotomy between IBD and FSGS studies in the literature, which impedes the simultaneous detection of IBD and FSGS, and, thereby, the inference of long- and short-term demographic signatures on spatial genetic structure. A spatially dense sampling of non-disturbed populations across the range would be necessary, however the sampling requirements for inferring FSGS at a given site can be an order of magnitude higher than that for IBD. Further, various ecological and disturbance factors may also affect the optimal size of the sampling area of FSGS studies (Vekemans & Hardy, 2004). Generally speaking, the sampling area should be as small as possible to capture only short-term processes, but it should also contain a sufficiently large number of individuals to precisely estimate the slope of the regression of kinship coefficients against geographic distances or the spatial autocorrelation of allele frequencies (Leblois et al., 2003; Smouse et al., 2008). The level of polymorphism in genetic markers can also vary across the species range, and influence the power of a data set for estimating kinship (Wang, 2016) or the spatial correlograms. The same genetic data that are used to estimate the spatial autocorrelation of allele frequencies may also be used to partially reconstruct the pedigree, which can be used to understand the mechanisms that maintain FSGS, such as the unequal reproductive success among individuals (e.g. Gerzabek et al., 2017; Oddou-Muratorio et al., 2018; Paluch et al., 2019) or sex specific dispersal dynamics (Aguillon et al., 2017).

Silver fir has recently attracted interest in the context of climate change for being more drought tolerant than other dominant forest tree species, such as Norway spruce (*Picea abies*) (Vitali et al., 2017; Vitasse et al., 2019). Fossil data also suggest that silver fir previously survived under much warmer than current temperatures, thus it could have become more adapted to drought (Tinner et al., 2013). There is little fossil evidence for the distribution of silver fir during the climatic oscillations of the Quaternary (past 2.5 Myr BP), but fossil records are common in the Holocene starting from approximately 11.6 kyr BP (Magri et al., 2017). The available data suggests that before the LGM (21 kyr BP) during warm glacial interstadials the species principally remained in refugia in the Italian and the Balkan peninsulas (Tinner et al., 2013). Two main maternal mtDNA lineages likely represent these two refugia (Liepelt et al., 2002). The range expansion from these refugia in southern Europe is currently ongoing, which is an exceptional event of the Quaternary history of the species (Willy Tinner, personal comm.). The southern part of the Carpathians was likely colonized from the Balkans while the eastern part from the Apennines (Liepelt et al., 2002, 2009; Gomory et al., 2012; Bosela et al., 2016). The Alps were most likely colonized from the Northern Apennines but two additional routes have likely contributed, namely from southern France and from the Balkans through the Adriatic coast (Burga et al., 2001; Terhürne-Berson et al., 2004; Liepelt et al., 2009; Cheddadi et al., 2014; Ruosch et al., 2016; Piotti et al., 2017). The Apennines hosted at least three refugial areas during the LGM, whose exact delimitation has been described by Piotti et al. (2017). Silver fir populations from the Pyrenees show a high genetic differentiation from the rest of species range, but belong to the western mtDNA haplotype group, suggesting that Pyrenees have also been colonized from the Apennines, but much earlier then the last glacial cycle (Liepelt et al., 2009).

The main objective of this study was to identify demographic and environmental drivers of FSGS across the species range of silver fir. We studied natural silver fir populations with no or close-to-nature management, so we assumed that differences in FSGS were due to demographic history and/or environmental constraints/adaptation. We assembled published single nucleotide polymorphism (SNP) maker data for adult trees (Brousseau et al., 2016; Roschanski et al., 2016; Heer et al., 2018) from eight sites sampled across the species range. In particular, the sites represented the two mtDNA haplotype groups and cover all recolonization routes from effective refugia (Liepelt et al., 2002 and 2009). Further, two populations from Central Apennines and the Pyrenees that have been isolated from other western populations before the Holocene were included (Terhürne-Berson et al., 2004). At each site, a pair of low and high elevation populations were sampled that most likely have been separated from one another at more recent time scales. This hierarchical sampling design allowed us to disentangle the effects of the distant past, i.e. the colonization of Europe from glacial refugia during the Holocene, and those of recent demographic events, such as possible colonization of high elevation habitats during the past few generations (Chauchard et al., 2010; Hernández et al., 2019; Vitasse et al., 2019). We developed a proxy for the effects of distant past using range-wide IBD patterns, and hypothesized that populations close to the supposed refugia are in migration-drift equilibrium and exhibit no FSGS, while populations in recently colonized areas of the species range show some degree of spatial structuring. With respect to high vs low elevation population pairs, we hypothesized that both recent demographic history and potential effects of environmental adaptation or habitat quality could contribute to differences in FGSG. Indeed, previous studies on some of these populations showed evidence for adaptation to elevation (Brousseau et al., 2016; Roschanski et al., 2016), and the effect of environmental factors on FSGS have been identified in other Italian silver fir populations (Mosca et al., 2018). Although the scale of our study allowed to disentangle demographic effects at different time scales, the sampling was performed by different research groups, which raised the question whether differences in FSGS across the 16 populations reflect not only biological patterns, but eventual sampling artifacts. To this end, we performed a resampling of our largest populations to assess how different spatial sampling configurations might have influenced our results. Finally, we addressed the question of the maintenance of FSGS and genetic diversity across the range. To this end, we estimated the strength of FSGS in seedlings, as well as the distribution of reproductive success in adult trees, and dispersal parameters, such as the selfing rate and the gametic gene flow. We hypothesized that seedlings would exhibit a stronger FSGS than adults because of limited seed dispersal, especially in recently colonized, non-equilibrium populations.

## Materials and Methods

### Data sets

We assembled a data set of 1368 adult trees from previously published data from eight different sites across the species distribution range (Figure 1a and Table 1; Issole (ISS), Lure (LUR), and Vesubie (VES) from Roschanski et al. (2016); Apennines (APE), Fagaras Mountains (FAG), Pyrenees (PYR), and Ventoux (VEN) from Brousseau et al. (2016), and Bavaria (BAV) from Heer et al. (2018)). Each site was sampled at high and low elevations that were subsequently denoted with _H and _L after the population names (Table 1). The number of genotyped adult trees per population ranged from 43 to 250 (Table 1, Figure S1). Additionally, at three sites (APE, BAV and FAG) we sampled and genotyped 540 seedlings with height < 1m and diameter at breast height < 4 cm (Table 1, Figure S1). Similar sampling configuration and area were used across all sites, but differences in the terrain and in the population density imposed some differences, especially in the spatial distribution of seedlings. Further, the sites LUR, ISS and VES were considered as replicates of VEN and sampling effort was less important (Table 1), but still a nearly exhaustive sampling of adult trees in a given area appeared suitable for inclusion in this study. As a result, the distance between the two farthest trees ranged from 68 m (ISS_H) to 1397 m (BAV_L) (Table 1 and Figure S1). Sampled trees were geo-referenced using a GPS device (Garmin Ltd.; Olathe, KS, USA in APE, BAV, ISS, LUR, PYR, VEN and VES; Trimble Juno T41/5 - Sunnyvale, CA 94085, USA in FAG) or compass and laser distance-meter (Leica Geosystems AG, St. Gallen, Switzerland in APE).

**Figure 1.**
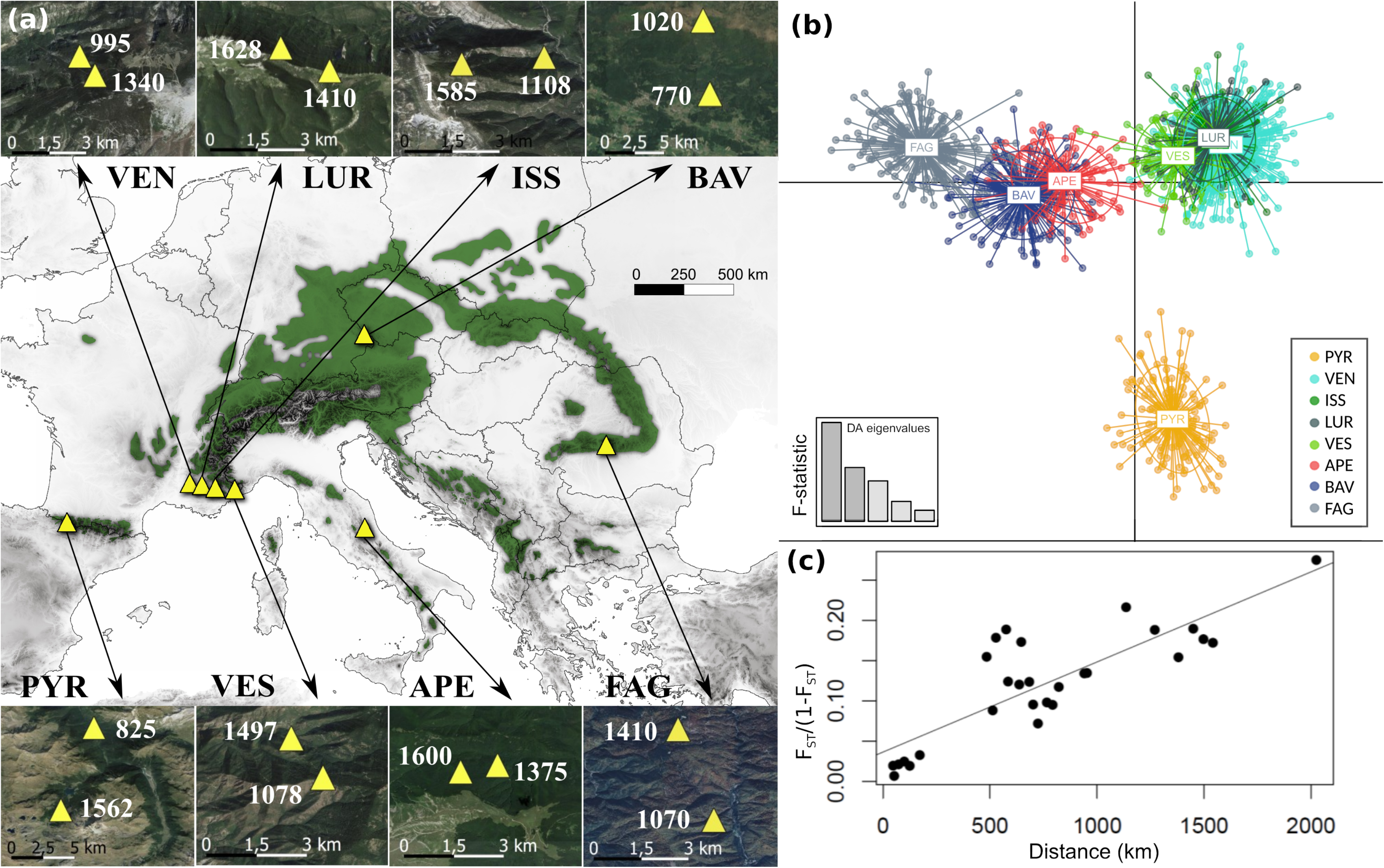
**(a)** Location of the study sites (main map, yellow triangles) and distribution of silver fir (*Abies alba* Mill.) (dark green). Small maps show the position of the high and low elevation populations withing each site. Site names are abbreviated using three letter codes; see Table 1 for full names. **(b)** Genetic clusters across the study sites inferred using a discriminant analysis of principal components (DACP) and the R package adegenet (Jombart, 2008). Clusters are represented in the space defined by the first two Linear Discriminant (LD) functions. **(c)** Isolation-by-distance in the studied populations.

**Table 1.** Geolocalization of the eight sampling sites, sample sizes, and the maximum distance between sampled adult trees in meters. At each site, a ow and a high population was sampled. At three sites seedlings were also sampled and their sampled sizes are in parenthesis.

| Site ID | Country | Site name | Latitude | Longitude | Elevation (m) |  | Sample size |  | Max distance (m) |  |
| --- | --- | --- | --- | --- | --- | --- | --- | --- | --- | --- |
|  |  |  |  |  | Low | High | Low | High | Low | High |
| PYR | France | Ossau Valley (Pyrenees) | 42.8550 | -0.4578 | 825 | 1562 | 81 | 82 | 198 | 444 |
| VEN | France | Ventoux (Alps) | 44.1812 | 5.27043 | 995 | 1340 | 214 | 250 | 242 | 187 |
| LUR | France | Lure (Alps) | 44.1142 | 5.83912 | 1410 | 1628 | 55 | 55 | 82 | 175 |
| ISS | France | Issole (Alps) | 44.0242 | 6.46244 | 1108 | 1585 | 49 | 47 | 92 | 68 |
| VES | France | Vesubie (Alps) | 43.9707 | 7.36577 | 1078 | 1497 | 43 | 45 | 121 | 62 |
| APE | Italy | Bosco Martese, Ceppo<br>(Central Apennine) | 42.6688 | 13.4322 | 1375 | 1600 | 46 (98) | 47 (57) | 198 | 234 |
| BAV | Germany | Bavarian Forest | 48.9468 | 13.4040 | 770 | 1120 | 100 (148) | 92 (143) | 1397 | 1031 |
| FAG | Romania | Arges (Fagaras Mountain) | 45.4411 | 24.6947 | 1070 | 1410 | 95 (46) | 94 (48) | 986 | 802 |

All individuals were genotyped using the same set of 267 SNP loci from 175 candidate genes using KASP assays manufactured by LGC Genomics (Middlesex, United Kingdom) (Roschanski et al., 2013 and 2016), with the exception of APE and FAG, where seedlings were genotyped at a subset of 211 loci. Among these, we only used loci with less than 15% missing data, a minor allele frequency > 5%, and that did not significantly deviate from the Hardy-Weinberg equilibrium in at least 75% of the populations (Table 2). Further, previous studies identified significant environmental and phenotypic associations at some of the loci (Brousseau et al., 2016; Roschanski et al., 2016; Heer et al., 2018). Here, we excluded only the outlier loci that coded for a non-synonymous amino acid change. This is because most of these outlier studies were under-powered due to a limited sample size, and, in fact, never identified the same set of loci (Csilléry et al., 2020). Further, even if some of the variants may code for phenotypic traits under selection, their effect sizes are likely too small to matter for estimating kinship at a short spatial scale. Taking together all of the above considerations, we could use a common set of 137 SNPs for the adult trees, and 116 SNPs for the seedlings.

**Table 2.**
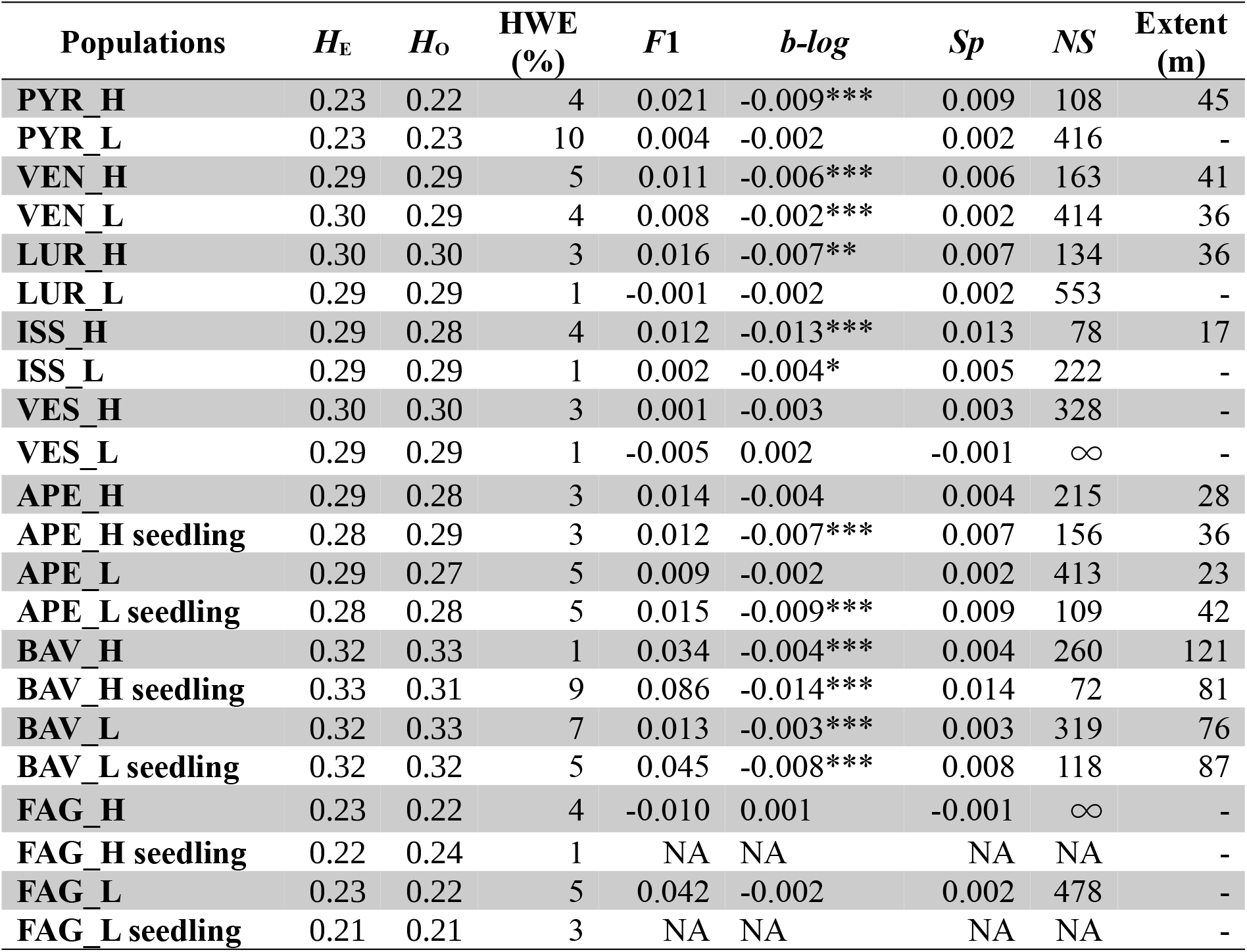
Genetic diversity and summaries of the Fine-scale Spatial Genetic Structure (FSGS) in silver fir across eight sites with a high (_H) and low (_L) elevation populations from each. Statistics are reported for adult trees, and, at three sites, for seedlings as well. Abbreviations: *H*E: Expected heterozygosity, *H*O: Observed heterozygosity, HWE: percentage of loci that significantly deviated from the Hardy-Weinberg Equilibrium (R package *pegas*, Paradis, 2010), *F*1: mean kinship of the first distance class (0-20 m), *b-log*: regression slope of kinship (*: p-value < 0.05, **: p-value < 0.01, ***: p-value < 0.001), *NS*: neighborhood size, NA: not available because the spatial clustering was too strong (see Materials and Methods for details), Extent (m) is the extent of FSGS in meters estimated as the distance where *F*ij was significant with the even sample sizes method.

### Environmental variables

The habitat at each site was characterized using a combination of climate, soil, and species distribution data. Downscaled historical climate data at 1 km resolution was obtained from the CHELSAcruts time-series (Karger et al., 2017; https://chelsa-climate.org/chelsacruts/) from 1901 to 1990. Monthly mean, minimum, and maximum temperatures and monthly precipitation were extracted and used to calculate the annual mean temperature, isothermality, and annual total precipitation using the R package *dismo* (Hijmans et al., 2017). We calculated only these climatic variables because they have been shown to drive adaptation in silver fir (e.g. Gazol et al., 2015; Csilléry et al., 2020), and we had to limit the number of variables to be tested because of the limited number of observations (16 populations). Soil available water capacity (AWC) was extracted at a 250 m resolution from the SoilGrids data set (Hengl et al., 2017). Measures of species composition were derived from the EU-Forest, which is a 1 km resolution tree occurrence database for Europe (Mauri et al., 2017). The occurrences of 14 tree species within areas of 0.3 radius circles (∼33 km) around each study populations were extracted.

### Statistical analyses

First, we assessed genetic diversity parameters and IBD patterns across the species range. We characterized genetic diversity and divergence at the level of sites and populations. Expected heterozygosity (*H*_E_) and observed heterozygosity (*H*_O_) were calculated for each site, population and cohort (i.e. adults and seedlings) using SpaGeDi 1.5b (Hardy & Vekemans, 2002), as well as pairwise *F*_ST_ among sites, between low and high elevation population pairs and between cohorts. Statistical significance of pairwise *F*_ST_ values was tested by 10,000 permutations. We performed a hierarchical analysis of molecular variance (AMOVA) to determine the partition of the genetic variation within and among sites and populations using the R package *pegas* (Paradis, 2010). We applied discriminant analysis of principal components (DACP) to identify genetic clusters (Jombart et al., 2010) implemented in the R package *adegenet* (Jombart, 2008). DACP identifies synthetic variables, so called Linear Discriminants (LD), that differentiate among a priori defined groups (here, populations), while minimizing variation within inferred genetic clusters. We used the function *Xval.dapc* to find the optimal number of principal components (PCs) to be retained, and *find.clusters* to find the optimal number of clusters based on the Bayesian Information Criterion (BIC). We used the number of populations (16) as the maximum number of clusters. The cross-validation procedure suggested that it was sufficient to include the first 120 PCs. Although the lowest BIC was observed with a grouping in nine clusters, the drop in BIC values was no longer marked after six clusters (Figure S2), where LUR, ISS, and VES belonged to the same “cluster 5” (Figure S2). Since DAPC can be sensitive to sampling density, we also carried out the DAPC analysis using only the two most distant French populations (VEN and VES). Finally, we plotted *F*_ST_/(1-*F*_ST_) against pairwise geographic distances among populations to illustrate the range-wide IBD pattern, but we did not perform a formal Mantel test because the spatial scale was several orders of magnitude larger than the dispersal distance of the species and because the studied populations experienced different long-term demographic dynamics and originated from different glacial refugia (Liepelt et al., 2002 and 2009).

Second, FSGS was characterized using two methods, the *Sp* statistic (Vekemans & Hardy, 2004) and multi-site spatial autocorrelation analyses (Smouse et al., 2008). *Sp* is estimated from a regression of pairwise kinship coefficients against the logarithm of pairwise spatial distances as -*b-log*/(1-*F*_1_), where *b-log* is the slope of the regression and *F*_1_ is the average kinship coefficient among individuals of the first distance class (Vekemans & Hardy, 2004). We estimated *Sp* for each population and each age cohort using the software SpaGeDi 1.5b. We used the pairwise kinship coefficient (*F_ij_*) of Loiselle et al. (1995), both with even distance classes (20 m) and even sample sizes per distance class. The spatial extent of FSGS was estimated as the distance where *F*_ij_ was significant with the even sample size method, as this option ensured that an adequate number of individuals were placed in each distance class. The statistical significance of *F*_ij_, *F*_1_, and *b-log* was tested using 10,000 permutations of individual locations among individuals. Multi-site spatial autocorrelation analyses of allele frequencies were performed using GenAlEx 6.5 (Peakall & Smouse, 2012) using even distance classes (20 m). The statistical significance of the autocorrelation was determined using 999 permutations by randomizing genotypes among distance classes. We also applied the non-parametric heterogeneity test described by Smouse et al. (2008) to test the statistical significance of the spatial autocorrelation patterns observed among populations.

Third, we tested the sensitivity of FSGS to sample size and sampling configuration using resampling. For this purpose, we chose VEN_H and VEN_L populations because of their high sample sizes (Table 1). The resampling was performed by applying three strategies: (1) resampling randomly within the sampling area, (2) reducing the size of the sampling area, and (3) resampling along linear transects (Figure 4). The first resampling approach was performed selecting 50 random individuals 20 times using R (R Core Team, 2019). The second resampling approach was carried out in the program QGIS (QGIS Development Team, 2016), where equal-sized (0.45 ha for VEN_H and 0.65 ha for VEN_L), choosing 20 circles with random centers placed within the central area of the population, so that the whole circle could always be inside the population. Finally, the third resampling approach required sampling across 20 linear transects, randomly drawn across each population using the R packages *sf* (Pebesma, 2018), *sp* (Pebesma & Bivand, 2005), and *raster* (Hijmans, 2015). The maximum and minimum coordinates within each population were used to create the first transect, and then the transect was rotated 20 times by 9°. FSGS analyses of the resampled data sets were carried out using an in-house python script that automated SpaGeDi, and using the even sample size option. We tested whether the resampling would have changed our conclusions in terms of the strength and extent of FSGS using one sample t-test, assuming that the estimates from the full data sets were the true values. Finally, we assessed the effect of sampling using all populations. For this, we calculated the correlation between summary statistics of FSGS (*F*_1_, *b*-*log*, *Sp*) and summaries of the sampling design, such as the area of the sampling site (m^2^), the maximum distance among the furthest individuals (m), and the density of sampling (number of samples/area of the sampling site) using R.

Fourth, we used a Structural Equation Modeling (SEM) approach to identify the potential demographic and environmental drivers of FSGS using the *lavaan* R package (Rosseel, 2011). Since we had only 16 populations, we were able to test only a limited number of variables. To partially circumvent this problem, we defined two models. In the first model, we tested the effect of demography. We included synthetic variables from the DAPC analyses (i.e. the first and second linear discriminant functions,LD1.DAPC and LD2.DAPC) as proxies for long-term demography, and elevation as a binary variable for short-term demography. In the second model, we tested the effect of the environmental variables. We simplified the occurrence data of 14 tree species using a Principal Component Analysis (PCA) using the *prcomp* function in R. The first three axes explained 78% of the variance of the original variables, thus we retained only these for the SEM analyses (hereafter PC1.spcomp, PC2.spcomp, and PC3.spcomp). PC1.spcomp was mainly determined by the presence of *Acer pseudoplatanus*, *Betula pendula*, *Fagus sylvatica*, *Pinus nigra*, *Quercus pubescens* and *Sorbus aucuparia*. PC2.spcomp was dominated by the presence of *Castanea sativa*, *Betula pendula*, *Picea abies*, *Pinus halepensis* and *Quercus ilex*, while PC3.spcomp was dominated by *Acer pseudoplatanus*, *Castanea sativa*, *Populus tremula* and *Quercus robur* (Figure S4). All variables except species composition were normalized using the *scale* function in R. For both models, we started with an initial model where several plausible functional relationships were hypothesized between *Sp* and explanatory variables (Table S7 and S8). Then, non-significant paths and variables were excluded step-by-step until the best explanatory model was found (Table S7 and S8). Based on Grace (2006) and Kline (2015), the best-fit model was selected using the following indices: Chi-square test (p-value > 0.05), the comparative fit index (CFI > 0.9), Bayesian information criterion, standardized root mean square residual (SRMR < 0.06) and root mean square error of approximation (RMSEA < 0.05).

Fifth, we tested for the presence and strength of FSGS in seedlings using the same method as described for the adult trees at the three sites where data for seedlings were available (APE, BAV, and FAG). We also tested for the heterogeneity in correlation of allele frequencies between cohorts (Smouse et al., 2008). Seedlings from FAG were excluded from these analyses because regeneration was limited to four patches in FAG_H and one patch in FAG_L (see Figure S1). In order to estimate dispersal parameters, we carried out parentage analysis. We used the likelihood-based approach implemented in the software CERVUS 3.0.7. (Kalinowski et al., 2007; Marshall et al., 1998). We used the “parent-pair” option with unknown sexes and the 95% or “strict” confidence levels. Simulation parameters were set as follows: simulation of 50,000 offspring, individual genotypes at a minimum of 68 loci, and a genotyping error rate of 0.01 (conform with the high precision of the KASP chemistry). The proportion of candidate parents included in the sample had a great influence on the proportion of assigned parent-offspring pairs. In order to choose the most probable value for each population, we performed the analysis with values of 50, 60, 70, 80, and 90% of candidate parents, and recorded the parent-offspring mismatch rates in each. For the final analysis, we used the highest proportion of candidate parents that gave the lowest mismatch rate across the tested values (Figure S5). We used the reconstructed genealogies of the offspring, together with the coordinates of both parents and offspring, to estimate (1) selfing rate, (2) mean variance and maximum reproductive success among all adult trees, defined as the number of gametes produced by each adult, (3) gametic gene flow rate, defined as the number of gametes that originated from outside the sampling area over the total number of gametes sampled, and (4) average parent-offspring distance. We also plotted and visually compared the distribution of individual reproductive success among populations.

## Results

### Genetic diversity and divergence across sites and populations

Genetic diversity was the highest in the Bavarian (BAV) populations (*H*_E_ = 0.32) and the lowest in the eastern and western range margin (FAG and PYR, *H*_E_ = 0.23, Table 2) populations. The hierarchical AMOVA revealed that most of the variance in allele frequencies was within populations (82.6%, *F*_ST_ = 0.174, p-value < 0.001, Table S1). However, a strong and significant genetic differentiation was detected among sites (14.8%, *F*_CT_ = 0.148, p-value < 0.001). In contrast, the differentiation between high and low elevation populations within sites was low, albeit significant (3.1%, *F*_SC_ = 0.0308, p-value < 0.001, Table S1). The most genetically differentiated pairs of sites were also the most geographically distant ones (Figure 1b and c, Table S2): PYR and FAG (*F*_ST_ = 0.275), and PYR and APE (*F*_ST_ = 0.216).

The DAPC analysis revealed that the first linear discriminant function (LD1) ordered the populations from east to west with the exception of PYR, which situated close to the French populations (VEN, LUR, ISS, VES) (Figure 1a). The second linear discriminant function (LD2) mainly separated the population from the Pyrenees (PYR) from the other populations (Figure 1b). The following linear discriminants appeared difficult to be interpreted (Figure S3). Although genetic clustering methods are known to be sensitive to sample size, the same clustering was obtained even after having excluded LUR and ISS, which balanced out the sample size (Figure S3). Finally, a rather clear IBD pattern was found across the species range (Figure 1c).

### FSGS across sites and populations in adult trees

We found a significant FSGS in eight out of 16 populations based on the *b-log* statistic (Table 2), and in seven populations based on the *r* statistic in the first distance class (0 - 20 m) (Figure 2). Based on the two tests taken together, populations from the sites BAV, ISS, and VEN, as well as LUR_H and PYR_H showed significant FSGS. Significant *Sp* values ranged from 0.003 to 0.013 (Table 2). The average extent of FSGS was 47 m (range: 17 m - 121 m) across all populations (Table 2). The spatial autocorrelation of allele frequencies did not differ globally among sites based on a test of heterogeneity (ω = 145, p-value = 0.64), while, in population-wise comparisons, APE_L differed significantly from several other populations based on the heterogeneity test (Table S4).

**Figure 2.**
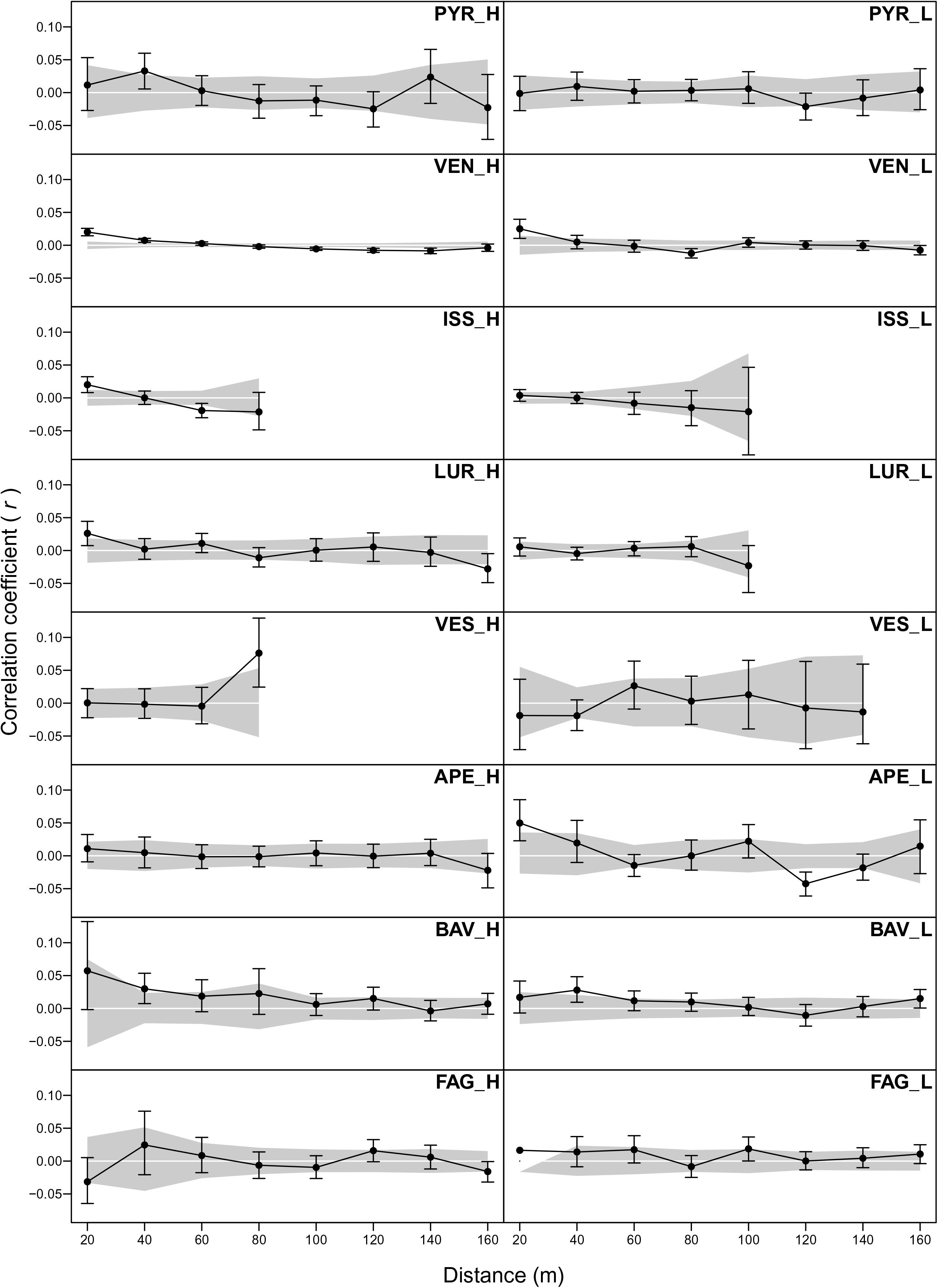
Fine-scale Spatial Genetic Structure (FSGS) across 16 silver fir (*Abies alba* Mill.) populations based on 137 SNPs using the software GenAlEx 6.5 (Peakall & Smouse, 2012). Correlation coefficients (*r*) are plotted against the maximum geographical distance between individuals in each distance class. Shaded areas represent the 95% confidence interval obtained through random shuffling of individual geographic locations (1000 times). Vertical bars around mean *r* values represent 95% confidence intervals generated by bootstrapping (10000 times) pair-wise comparisons within each distance class. Note that in FAG_L we could not estimate the confidence intervals in the first distance class due to the limited number of individuals.

### The effect of sampling scheme on FSGS

With all three methods of resampling, we obtained FSGS correlograms similar to those based on the full data set (Figure 4). Nevertheless, the *Sp* values based on a reduced number of samples did not always reflect the significant *Sp* values observed using the full data set, both in VEN_L and VEN_H (Table 2). While reduced area and linear transect resampling resulted in significant *Sp* values in 83% and 88% of the total number of resampling, respectively, random resampling showed significant FSGS only in 46% of the cases. In terms of the absolute value of *Sp*, all resampling methods provided consistent results with those obtained using the full data set in VEN_H (Figure 4, one-sample t-tests, p-value > 0.76; Table S6). In contrast, in VEN_L only random resampling method provided the same results based on the full data set (one-sample t-tests, p-value = 0.64), while both reduced area and linear transect resampling overestimated the *Sp* values (Figure 4, p-value < 0.001; Table S6). Finally, no correlation was found between the observed *F*_ij_, *F*_1_, *b*-*log,* or *Sp* values in the 16 populations and the area of the sampling site, the maximum distance among individuals, and the density of sampling (Pearson correlation < 1.59, p-value > 0.13).

### Demographic and environmental drivers of FSGS in adult trees

The best SEM model including demographic variables explained 47% of the variation in *Sp* values across the 16 populations (Table S7). *Sp* was significantly affected by elevation (High/Low), with high elevation populations exhibiting higher *Sp* values, and by the first discriminant function (LD1.DACP), with eastern populations having a stronger FSGS than western ones (Figure 4a). The best environmental SEM model explained 58% of the variation in *Sp* values (Table S8). The environmental variables that influenced *Sp* the most were elevation and isothermality (Figure 4b). Both effects were positive, thus FSGS was stronger in populations at higher elevations and where the diurnal temperatures oscillations were larger compared to seasonal variations. Although with weaker effects, AWC and species composition (PC2.spcomp) also influenced *Sp* values. Species composition itself was impacted by elevation, AWC, temperature and precipitation, thus these variables also had an indirect effect on FSGS (Figure 4b).

### Genetic diversity and FSGS in seedlings and dispersal parameters

Genetic diversity in seedlings was overall similar to that observed in adult trees (Table 2). We found weak, but significant genetic divergence between adult and seedling cohorts in APE and FAG, with *F*_ST_ values ranging from 0.002 to 0.007 (Table S3), but not in BAV. FSGS in the seedling cohort was significant in all populations (APE_H, APE_L, BAV_H, BAV_L), based on both the *b-log* and *r* statistics (Figure 5, Table 2). Overall, FSGS in seedlings (average *Sp* = 0.01, Table 2) was stronger than in adults (Two Sample t-test, t(18) = −2.87, p-value = 0.01). In contrast, the heterogeneity test of spatial autocorrelation between adult and seedling cohorts was not significant in any of the populations (Table S5). Finally, the extent of FSGS in seedlings ranged between 36 and 87 m (average of 62 m), thus slightly higher than in adults.

Parentage analysis revealed a strong reproductive skew in all six populations (Table 3, Figure 6). Only 54%, 25% and 6% of the adults in BAV, APE, and FAG contributed to the generation of sampled seedlings, respectively. The maximum individual reproductive success was recorded in APE_L, with 18 offspring assigned to a single adult tree (Table 3). The variance in individual reproductive success was similar in BAV_H and BAV_L (1.8) and APE_L (1.7), while lower in APE_H (0.5) and the lowest in FAG_H and FAG_L (0.1). Self-pollination occurred in APE_L and BAV_H and BAV_L, and was as high as 9% in BAV_H. The gametic gene flow rates were the highest in FAG, 87 and 95% in FAG_H and _L, respectively, because we could not identify the parents of most seedlings, and the lowest in BAV_H and BAV_L, but still 41 and 40%, respectively (Table 3). The mean parent-offspring distance varied among the three sites (APE: 40 m, BAV: 72 m, FAG: 130 m), but it was similar between low and high populations (Table 3), and comparable to the extent of FSGS estimated from the adult and seedling data (Table 2).

**Table 3.** Summaries of the parentage assignment per population using CERVUS 3.0.7. (Kalinowski et al., 2007; Marshall et al., 1998). The mean reproductive success is the mean number of offspring per adult tree. The variance in reproductive success is the variance of the number of offspring across all adult trees. The gametic gene flow rate was estimated as the number of gametes that originated from outside the sampling area (i.e. number of gametes for which no parent was found within the population) over the total number of gametes sampled (i.e. twice the number of seedlings) (Valbuena-Carabaña et al., 2005).

| Summary level | Dispersal parameter | APE<br>H | APE<br>L | BAV<br>H | BAV<br>L | FAG<br>H | FAG<br>L |
| --- | --- | --- | --- | --- | --- | --- | --- |
| Seedlings | Both parents identified (trios) (%) | 8 | 18 | 35 | 33 | 0 | 0 |
|  | One parent identified (%) | 28 | 43 | 48 | 54 | 10 | 26 |
|  | Self pollination (%) | 0 | 3 | 9 | 4 | 0 | 0 |
| Adults | With offspring (parents) (%) | 23 | 26 | 49 | 59 | 3 | 8 |
|  | Mean reproductive success | 0.5 | 1.7 | 1.8 | 1.8 | 0.1 | 0.1 |
|  | Variance in reproductive success | 3.0 | 18.4 | 7.3 | 5.5 | 0.1 | 0.2 |
|  | Maximum reproductive success | 11 | 18 | 11 | 11 | 3 | 3 |
| Populations | Gametic gene flow rate | 0.77 | 0.60 | 0.41 | 0.40 | 0.95 | 0.87 |
|  | Mean parent-offspring distance (m) | 44 | 35 | 77 | 66 | 119 | 141 |

## Discussion

### Genetic diversity of silver fir populations across the species range

Consistently with its large effective population size, *N_e_,* and its wind pollinated outcrossing mating system (Ennos, 2001; Vekemans & Hardy, 2004; Hardy et al., 2005), we found high levels of genetic diversity across the whole range of silver fir (average *H_E_* of 0.3; see Table 2). As changes in genetic diversity scale with 2*N_e_*, high levels of genetic diversity most likely still reflect the diversity of refugial populations during the Last Glacial Maximum (LGM; 21,000 years BP; Petit & Hampe, 2006). Populations from the French Mediterranean Alps (VEN, LUR, ISS, VES) that have been colonized from main effective refugium situated in the northern Apennines exhibited high levels of diversity (Table 2). The population from the central Apennines, APE, also exhibited a relatively high level of diversity (Table 2). This region likley represents a separate, smaller refugium that evolved independently from populations in the northern and southern Apennines (Piotti et al., 2017). In our study, APE exhibited a stronger similarity to the population from the Carpathians, FAG, than to those from the French Mediterranean Alps (Figure 1b), which is in agreement with some other studies (Brousseau et al., 2016; Martínez-Sancho et al. 2021), but our scattered sampling is clearly not sutable to refine the demographic history of silver fir. Genetic diversity was the lowest in the westernmost (Pyrenees, PYR) and the easternmost (Carpathians, FAG) extremes of the distribution range (Figure 1 and Table 1). Silver fir populations from the Pyrenees likely originated from the refugium in the northern Apennines as well, but they have become isolated well before the Holocene, as suggested by the presence of macroremains older than 15 kyr BP (Terhürne-Berson et al., 2004). The Carpathians have been colonized during the Holocene from a relatively small refugium situated on the Balkan (Gömöry et al., 2004, 2012; Liepelt et al., 2010; Tinner et al., 2013), which is in agreement with the relatively low levels of genetic diversity in FAG. In this study, the highest diversity was observed at the German site, BAV, situated north of the Alps. These populations might have also been colonization from the northern Apennines but via another colonization route through the Dinaric Alps (Liepelt et al., 2009). Here, we found that BAV clustered together with FAG and APE (Figure 1b), thus we may speculate that different lineages might have admixed north of the Alps leading to an elevated level of diversity (Petit et al., 2008).

### FSGS across the species range and its determinants

We found that silver fir was characterized by a moderate and significant FSGS at most parts of the species range (Figure 2, Table 2). Among the previous studies of FSGS in silver fir, Sagnard et al. (2011) and Leonarduzzi et al. (2016) found a weak or absent FSGS in French and Central Italian populations, respectively, while a relatively strong FSGS was reported from populations from the Italian Alps (Mosca et al., 2018) and the Western Carpathians (Paluch et al., 2019). Based on our results together with the above mentioned studies, *Sp* in silver fir ranges from 0 to 0.013 with a global mean *Sp* of 0.005 (CI95% [-0.001, 0.01]). Silver fir is characterized by a long generation time, and heavy seeds (Royal Botanic Gardens Kew, 2015) and pollen grains (Eisenhut, 1961), which could contribute to a limited dispersal (Amm et al., 2012), and to the maintenance of FSGS across generations. Further, silver fir often occurs in association with other species such as Norway spruce and European beech, which causes the isolation from conspecifics and the clustering of relatives (Aussenac, 2002; Dobrowolska et al., 2017), thus the maintenance of FSGS.

In this study, we identified both demographic and environmental drivers of FSGS across the species range. Among the demographic proxies, not surprisingly, the most recent demographic event, i.e. the supposed colonization of high elevation habitats, left the strongest signal on FSGS (Figure 4a). Nevertheless, the first discriminant function, ordering populations along an east-west biogeographical gradient, also left a detectable signature on FSGS (Figure 4a), suggesting that FSGS may conserve the signature of demographic events on deep demographic time scales, potentially reaching back to the LGM. These results confirm that demographically younger populations are more likely to exhibit a spatial genetic structuring, i.e. higher elevation and, to some extent, eastern populations were more likely to exhibit a stronger FSGS. Several previous studies have already demonstrated that recently colonized populations exhibit a stronger spatial structure than core populations (e.g. Gapare & Aitken, 2005; De-Lucas et al., 2009; Pandey & Rajora, 2012). This phenomenon can be mediated by stand density and its effect on the strength of local genetic drift (Hamrick et al., 1993; Young & Merriam, 1994). Our results are also concordant with independent evidence about recent range expansion of the species during the past decades (Vitasse et al., 2019). While silver fir has been declining in warm and dry areas, especially at the limit of its distribution range and in low elevations and in the Mediterranean (e.g. Gazol et al. 2015; Dobrowolska et al., 2017; Hernández et al., 2019), the species has been reported to expand towards high elevations across the whole range (Chauchard et al. 2010; Hernández et al., 2019; Vitasse et al., 2019). This process was facilitated by the reintroduction of large predators that reduce ungulate browsing pressure on silver fir (Apollonio et al., 2010; Chauchard et al., 2010), and this might open new habitats suitable for the species.

In this study, we also searched for environmental drivers of FSGS and found that the model including habitat variables explained 11% more variance in FSGS that the model including only the three demographic proxies (Figure 4a). In contrast, we found that, among habitat variables as well, elevation was the most important driver of *Sp*, more important than climate or species composition (Figure 4b). Although these results emphasize that differences in *Sp* values were more likely reflecting demographic effects than differences in the environmental conditions, high elevation populations can also be characterized by harsher environmental conditions, thus we could not exclude that climatic constrains or adaptation might also have contributed to the higher FSGS observed at high elevations. For example, many of our high elevation sites had a high snow cover in winter and were situated on scree slopes (e.g. VEN, LUR, ISS). We also found a positive effect of isothermality, with larger variances in temperatures associated with a higher FSGS. The other climatic variables had an indirect effect on FSGS, through their influence on species composition (Figure 4b). We note that high isothermality was a good indicator of continental climate, thus this effect could also simply coincide with the east-west demographic axis. Mosca et al. (2018) studied the role of climatic variables on FSGS in silver fir, and found a weak negative correlation between FSGS and spring precipitation in the Italian Alps. At a local scale, different climatic factors could indeed play a role in the strength of FSGS. However, regional studies can be limited in revealing the drivers of FSGS because the environment of each population is only compared to similar populations from the same region.

### Limitations of this study: the effect of sampling

Sampling is a key issue in all IBD and FSGS studies, where separating biological patterns from sampling artifacts can be challenging (Vekemans & Hardy, 2004; Smouse et al., 2008; Legendre & Fortin, 2010; Meirmans, 2012). This study had to face this challenge in particular because it was aimed at understanding both range-scale and population-scale processes, as well as their interaction. At the range-scale, although our sites represented most genetic lineages, several parts of the range were underrepresented (Figure 1). Thus, it is possible that we did not have sufficient power to tease apart the effects of demography and the environment. Measures of FSGS are also sensitive to several aspects of sampling, such as sampling area, configuration, and density, which makes among-population comparisons particularly challenging (Vekemans & Hardy, 2004; Smouse et al., 2008). Although the global test of heterogeneity in spatial autocorrelation of allele frequencies did not show evidence for variation in FSGS across the range, we argue that this result appeared rather conservative given the strong population differences in *Sp* values and their clear link to demography (Table 2, Figure 4). Unfortunately, little information is available about the performance of this test across a large number of populations. We found that a resampling experiment on our own data can be recommended for understanding the limitations of a data set. The main conclusion from our experiment was that, when tree density was low (VEN_L), the reduced area or linear transect sampling lead to an overestimation of *Sp* values (Figure 3b and c). In contrast, the FSGS analyses appeared robust to any kind of resampling when there was a high tree density (VEN_H, Figure 3). Nevertheless, when we considered the observed data from all 16 populations, we found no evidence for a consistently higher *Sp* in populations where the sampling area was reduced, such as LUR, ISS or VES, and we found a strong and significant FSGS in BAV, where the sampling density was low (Figure S1, Table 2, Table S6). Finally, we acknowledge that the sampling error of *Sp* can be very high (see error bars on Figure 3), so we reiterate the warnings of previous studies that large sample size and several populations across the range are needed to draw robust conclusions (Vekemans & Hardy, 2004; Smouse et al., 2008).

**Figure 3.**
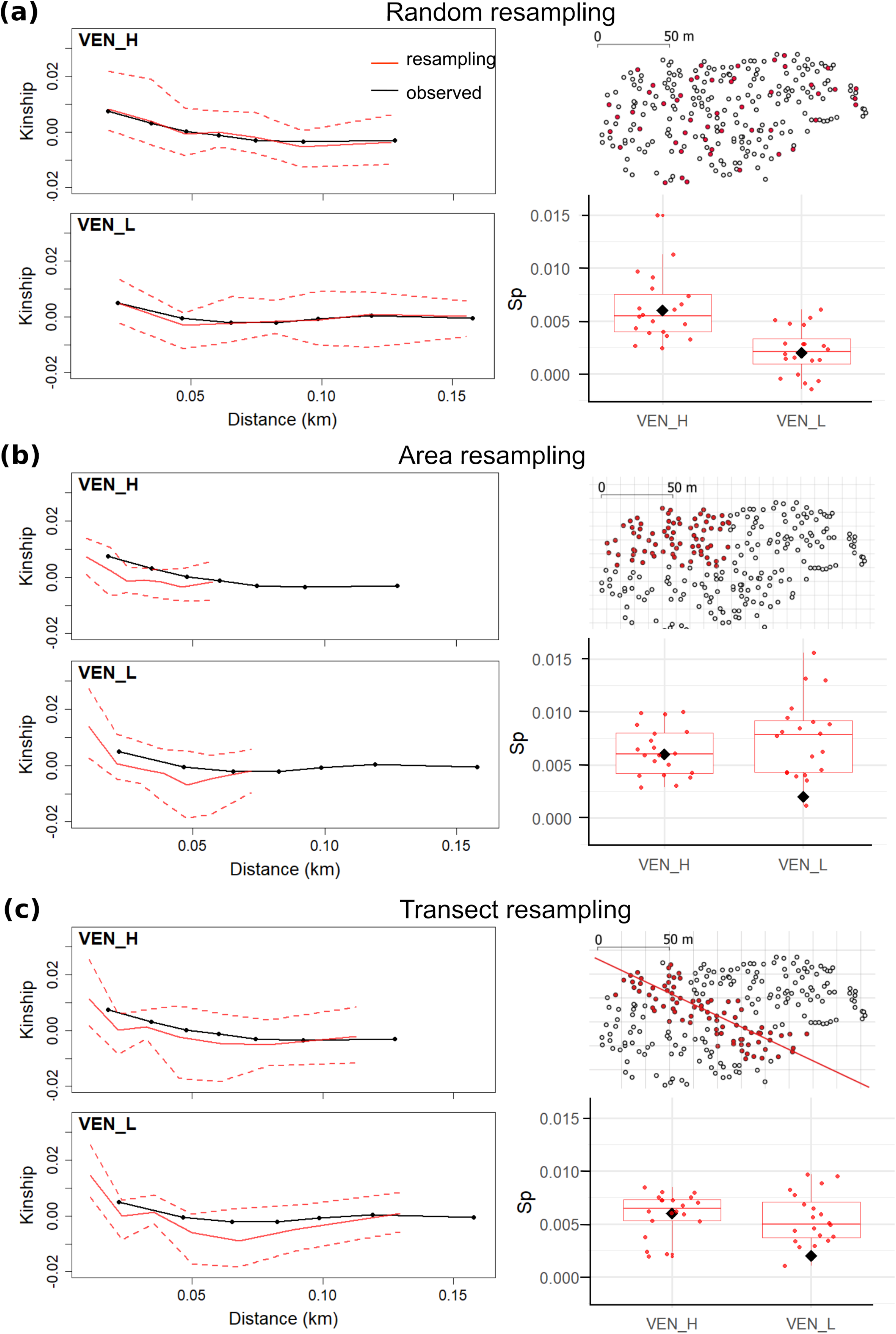
Assessment of the sampling configuration on inferences about the Fine-scale Spatial Genetic Structure (FSGS) using two silver fir (*Abies alba* Mill.) populations, VEN_H and VEN_L, and the software SpaGeDi 1.5b (Hardy & Vekemans, 2002). Three spatial resampling methods have been tested: random resampling **(a)**, reduced area **(b)**, and linear transect **(c)**. For each of the three methods: Maps illustrate the resampling method and correlograms show the average kinship coefficients plotted against the mean distance between individuals. Full lines indicate the mean of 20 resamplings and dotted lines indicate the lower and higher limits of kinship values. Box plots show the distribution of *Sp* statistics after resampling. Black diamonds on boxplots illustrate the original *Sp* of all samples.

**Figure 4.**
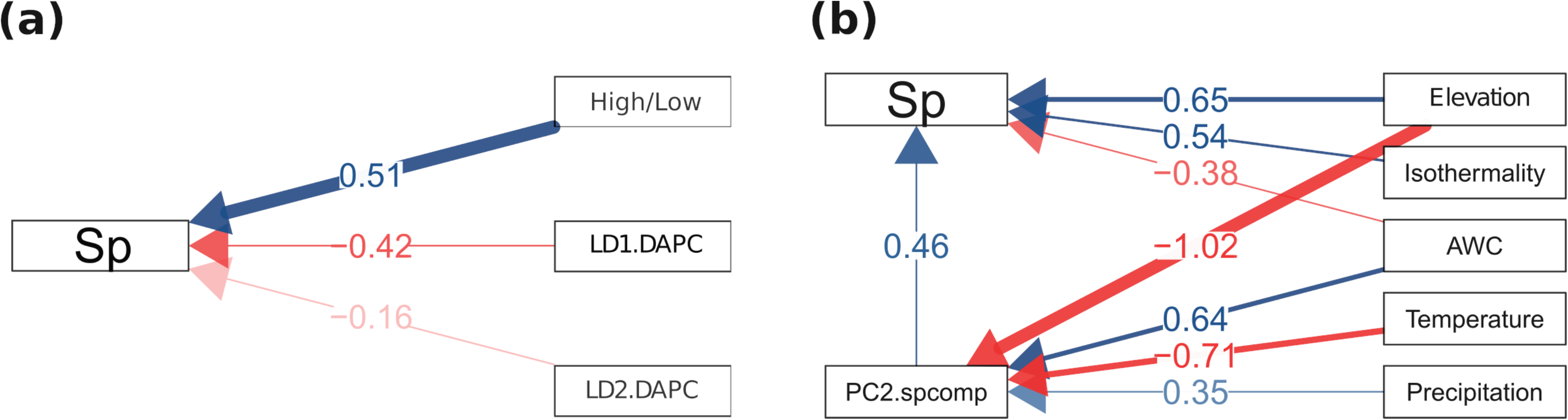
Path diagram of the two final Structural Equation Models (SEM). **(a)** SEM estimating the causal effects of demography including High/Low status and the first two linear discriminants (LDs) from the DAPC analysis on *Sp*. **(b)** SEM estimating the causal effects of Temperature, Isothermality, Precipitation, soil available water capacity (AWC) and elevation on *Sp* and on species composition (PC2.spcomp) synthetised as the second principal component of all species composition variables. We also hypothesized an indirect effect of the explanatory variables on *Sp* via PC2.spcomp. Lines in green correspond to a positive effect (or a positive correlation between variables) whereas lines in red correspond to a negative effect or correlation. Color intensity is positively related to the significance of the relationship between two variables. Standardized path coefficients between variables are indicated on the arrows.

### The maintenance of FSGS across generations

In small and isolated populations, elevated inbreeding levels might reduce fitness of subsequent generations and threaten forest resilience in the long term (Restoux et al., 2008; González-Díaz et al., 2012). Comparing FSGS between life stages (e.g. adult and juvenile cohorts) may identify when populations have undergone disturbances and help understand the intrinsic mechanisms underlying patterns of FSGS (Segelbacher et al., 2010). Here, we found that *Sp* was significantly stronger in the seedling cohorts compared to adults (Figure 2 and 5, Table 2). This result likely reflects seed shadows around adult trees (González-Martínez et al., 2002; Ng et al., 2004; Troupin et al., 2006). Parentage analyses also supported the existence of seed shadows, especially at high elevations: estimates of the gametic gene flow were consistently higher in the higher elevation populations than in the lower ones (Table 3). This result is in line with our hypothesis that higher elevation populations were more recently colonized, but may also suggest populations at low or intermediate elevations act as continuous sources of gene flow for higher elevation populations (Herrera & Bazaga, 2008). In BAV, seedlings mainly originated from parent trees within the sampling area and the proportion of selfing was also high (Table 3). This result could reflect the low silver fir density in Bavaria, where the proportion of Norway spruce (*Picea abies* K.) is the highest among all studied sites. In contrast, at the high density APE site, regeneration to a large extent relied on pollen and seed immigration (Table 3). A similar effect of stand density on the outcrossing rate has been documented in Mediterranean silver fir stands (Restoux et al., 2008). In FAG, even though seedlings occurred in spatially aggregated regeneration spots, we were able to identify the parents of only a few seedlings (Table 3). Since we cannot exclude the possibility that parent trees have been cut, estimates of the dispersal parameters from FAG should be treated with caution.

**Figure 5.**
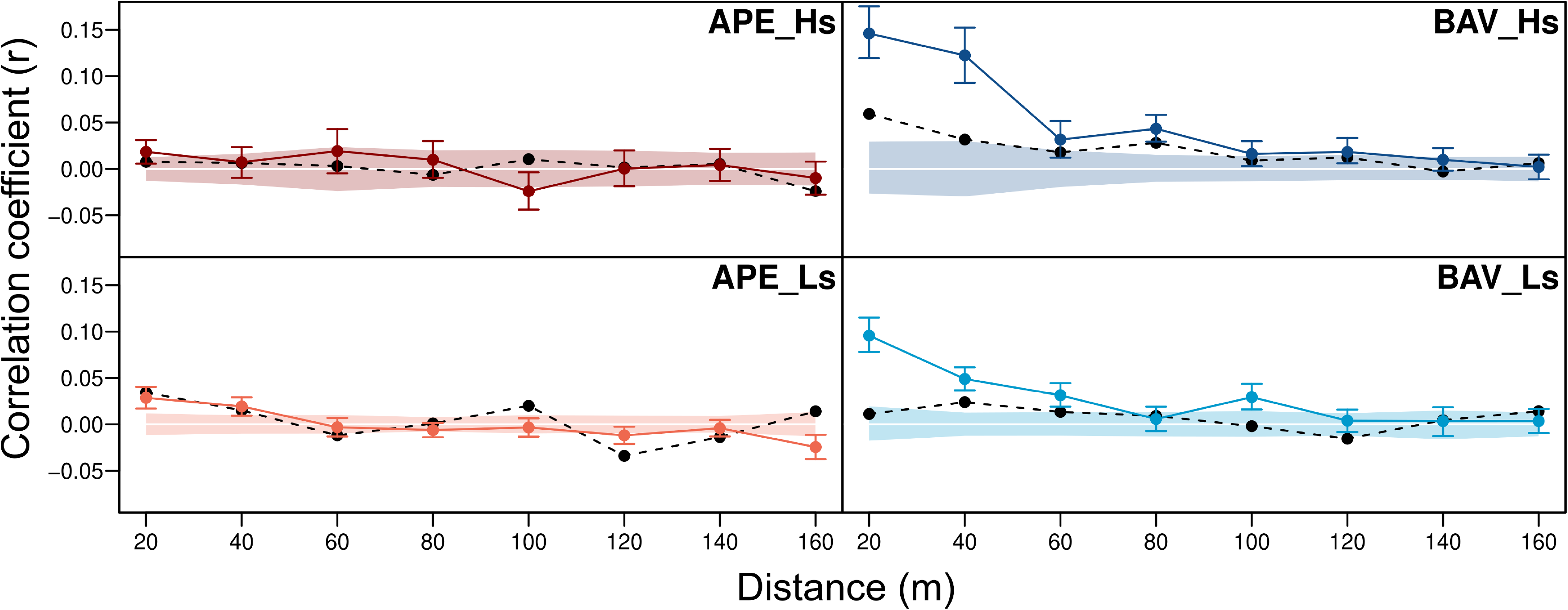
Fine-scale Spatial Genetic Structure (FSGS) across four silver fir (*Abies alba* Mill.) population pairs, comprising adult and seedling cohorts, using the software GenAlEx 6.5 (Peakall & Smouse, 2012). Correlation coefficients (*r*) were plotted against the maximum geographical distance between individuals in each distance class. Full lines refer to autocorrelation analyses of seedlings and dashed lines refer to autocorrelation analyses of adult trees.

The average number of seedlings per adult tree at our sites varied between 0.1 and 1.8 seedlings, with a single tree having 18 seedlings at the most extreme case in APE (Figure 6). Such pronounced reproductive skew is commonly observed in trees (e.g., Chybicki & Burczyk, 2013; Oddou-Muratorio et al., 2018), and can be driven by genetic or micro-environmental differences. For example, Avanzi et al. (2020) showed that in Norway spruce, trees that were able to maintain high-growth rates despite environmental growth limitations had the highest reproductive success. The assessment of reproductive success also greatly depends on the ontological stage. Indeed, it is well known that in perennials, the shape of the size frequency distribution changes greatly in populations through time, particularly when the density is high (e.g., Dodd & Silvertown, 2000; Herrera & Jovani, 2010). Gerzabek et al. (2017) studied maternal reproductive success across time in a oak stand, and found a marked inequality in reproductive success at the time of seedling emergence, that was considerably reshuffled by later seedling mortality. In our study, not all age classes were present at all sites, suggesting that the date of mast years and selective seedling mortality could have contributed to differences in the estimated reproductive success among sites (Davi et al., 2016). Overall it appears from our study that estimates of dispersal parameters, and also of FSGS in seedlings, are not directly informative for the FSGS of the next generation of adult trees.

**Figure 6.**
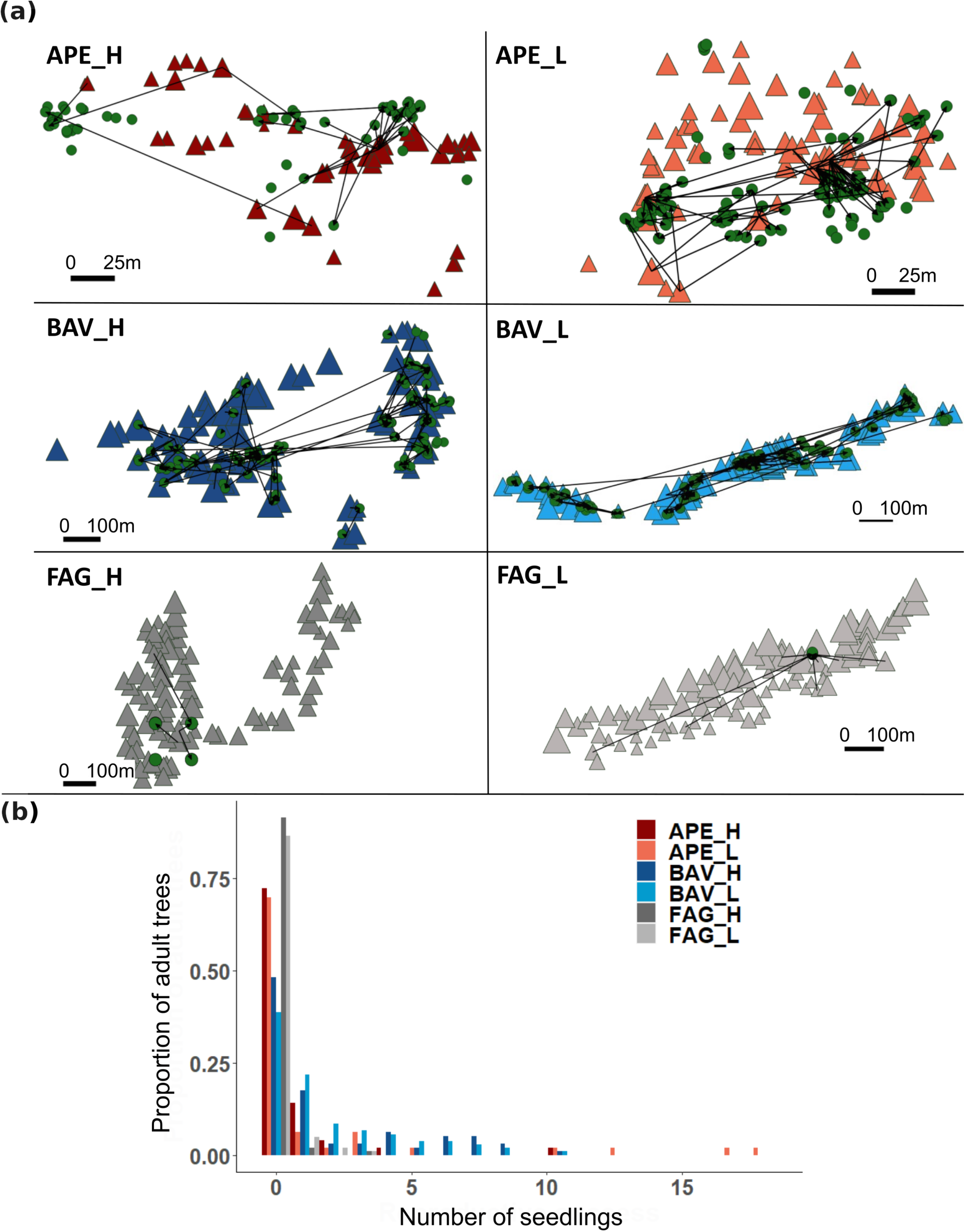
**(a)** Maps of all sampled adults (triangles with size proportional to the diameter at breast height) and seedlings (dots) and parent-offspring relationships (black arrows) inferred using CERVUS 3.0.7. (Kalinowski et al., 2007; Marshall et al., 1998) at three silver fir (*Abies alba* Mill.) sampling sites, each represented by its high (_H) and low (_L) elevation populations. **(b)** The distribution of reproductive success (i.e. the number identified seedling per adult tree within populations) in the same six populations with matching colour codes.

### FSGS as an indicator for monitoring genetic diversity?

Over the past decade, several studies and reviews have documented the apparent resilience of widespread tree species to the genetic consequences of habitat fragmentation and disturbance (Kramer et al., 2008; Bacles & Jump, 2011). Indeed, forest trees have several mechanisms to overcome genetic diversity loss and fitness reduction. For example, long distance pollen dispersal may assure connectivity among populations (Kremer et al., 2012), or overlapping generations could slow down the loss of genetic diversity (Austerlitz et al., 2000). While in an idealized Wright–Fisher population the amount of neutral genetic diversity is determined by the effective population size, *N_e_*, and its historical fluctuations (Ellegren & Galtier, 2016), in outcrossing wind-pollinated species with large *N_e_*, such as many forest trees, monitoring the average genetic diversity can be uninformative because changes in the genetic diversity scale with 2*N_e_*. It is often argued that populations with a higher genetic diversity may resist better or recover more quickly from a disturbance (Sgrò et al., 2011; Wernberg et al., 2018), and the average genetic diversity (Porth & El-Kassaby, 2014) or contemporary *Ne* (Hoban et al., 2020; Hoban et al., 2021) are commonly proposed indices for diversity monitoring (but see also a critique by Fady & Bozzano, 2020). In this study, we found no difference in the average genetic diversity between low and high populations, but we did in terms of FSGS. Other authors have also reported that forest management and disturbance regimes left a signature on FSGS but not on the average level of genetic diversity (Paffetti et al., 2012; Piotti et al., 2013; Leclerc et al., 2015; Sjölund & Jump, 2015). The analysis of the spatial distribution of genetic diversity, i.e. FSGS, allows to estimate the local equivalent of *N_e_*, the neighborhood size (*NS)* (Wright, 1943) as 1/*Sp* (Table 2). *NS* might be a useful indicator for monitoring diversity in forest trees as it is potentially sensitive to recent demographic changes and disturbances.

## Acknowledgements

EIM is supported by the Hungarian State doctoral program implemented at the Hungarian University of Agronomy and Life Sciences, School of Horticultural Science. KC was supported by a Swiss National Science Foundation grant (CRSK-3_190288) while working on this manuscript. CA, AP and GGV have been partially supported by resources available from the Italian Ministry of University and Research (FOE-2019) under the project “Climate Change” (CNR DTA.AD003.474). For genotyping the seedlings, the authors acknowledge the financial support of the ERAnet BiodivERsA project TipTree (ANR-12-EBID-0003 granted to LO, BF and GGV) and the German Federal Ministry of Education and Research (01LC1202A granted to LO). For field and lab work in Romania we acknowledge the National Research Program NUCLEU, Romania Ministry of Education and Research. We thank the Bavarian Forest National Park for supporting the fieldwork in Germany; the team of the INRAE UEFM in Avignon for sample collection and Anne Roig for lab work in France; the Gran Sasso e Monti della Laga National Park for logistic support, Cristina Leonarduzzi for help during the field work, and Catia Boggi and Ilaria Spanu for technical assistance and lab work in Italy; the team of INCDS Simeria for fieldwork in Romania and Ana Iordan for lab work.

## Author contributions

KC, MH, BF and GGV designed the study. KC, BF, GGV, and LO raised funds. LO and KH collected the seedling data in Germany (site BAV), AP and CA in Italy (site APE), and DP and FP in Romania (site FAG). EIM performed all analysis with help from KC. EIM, MH and KC wrote the first draft, and all authors contributed to improve the final manuscript.

## Data availability statement

New geo-referenced seedling SNP data associated with this manuscript will be available on Dryad upon acceptance. Different versions of the adult SNP data had already been published, see Materials and Methods for references, nevertheless they will also be made available upon acceptance using our filtering.

