## Supplementary Materials for "Fine-scale spatial genetic structure across the species range reflects recent colonization of high elevation habitats in silver fir (*Abies alba* Mill.)"

MOLECULAR ECOLOGY

### Table of Contents:


| | |
|---|---|
| **Table S1**. AMOVA of the 16 adult silver fir (*Abies alba* Mill.) populations. | Page 2 |
| **Table S2**. $F_{ST}$ among sites and pairwise $F_{ST}$ between high and low elevation population pairs. | Page 3 |
| **Table S3**. $F_{ST}$ among adult and seedling cohorts across six populations, and between low and high elevation population pairs in seedlings. | Page 4 |
| **Table S4**. Multi-class test statistic ($\omega$) and p-values for each pair of populations. | Page 5 |
| **Table S5**. Multi-class test statistic ($\omega$) and p-values between adults and seedlings. | Page 6 |
| **Table S6**. Sensitivity of *Sp* to the spatial configuration of the sampling in the two VEN populations, and with three resampling methods. | Page 7 |
| **Table S7**. Result of the structural equation modelling (SEM) analyses of *Sp* and demography with the best model. | Page 8 |
| **Table S8.** Result of the structural equation modelling (SEM) analyses of *Sp* and environmental variables with the best model. | Page 9 |
| **Figure S1**. Location of each sampled tree in each population separately. | Page 11 |
| **Figure S2**. Discriminant analysis of principal components (DAPC) results.: (A) BIC values versus number of clusters and (B) clusters versus populations. | Page 12 |
| **Figure S3.** Population structure revealed by the discriminant analysis of principal components (DAPC): Scatterplots based on the DAPC output for six assigned genetic clusters. | Page 13 |
| **Figure S4.** Situation of the 16 population locations along the first four Principal Components of species composition. | Page 14 |
| **Figure S5**. Representation of mismatch rates with different proportions of sampled parents (50. 60. 70. 80 and 90 %) used in the simulations for the parentage analyses. | Page 15 |

MOLECULAR ECOLOGY

**Table S1:** Analyses of molecular variance (AMOVA) of the 16 adult silver fir (*Abies alba* Mill.) populations using the R package *poppr* and the method *pegas*. P-values are based on 10000 permutations.

| Source of Variance | df | Sum Squares | Variance components | % |
|---|---|---|---|---|
| **Sites** | 7 | 6379.447 | 5.085 | <0.001 |
| **Elevation** | 8 | 841.917 | 0.902 | <0.001 |
| **Error** | 1352 | 38328.598 | 28.350 | |
| **Total** | 1367 | 45545.961 | | |

| Level of hierarchy | Fixation indexes |
|---|---|
| **Amog sites** | $F_{CT}$=0.148 |
| **Between elevations within sites** | $F_{SC}$=0.031 |
| **Among populations** | $F_{ST}$=0.174 |

MOLECULAR ECOLOGY

**Table S2.** $F_{ST}$ among sites and between high and low elevation population pairs using SPAGeDi. *: p-value < 0.01 from 10000 permutations.

| | $F_{ST}$ among sites | | | | | | | $F_{ST}$ |
|---|---|---|---|---|---|---|---|---|
| | **ISS** | **VES** | **LUR** | **VEN** | **BAV** | **APE** | **PYR** | **high-low** |
| **ISS** | | | | | | | | 0.022* |
| **VES** | 0.021* | | | | | | | 0.005* |
| **LUR** | 0.007* | 0.019* | | | | | | 0.024* |
| **VEN** | 0.025* | 0.033* | 0.019* | | | | | 0.016* |
| **BAV** | 0.098* | 0.072* | 0.095* | 0.117* | | | | 0.005* |
| **APE** | 0.124* | 0.088* | 0.120* | 0.123* | 0.095* | | | 0.001 |
| **PYR** | 0.188* | 0.173* | 0.178* | 0.155* | 0.188* | 0.216* | | 0.020* |
| **FAG** | 0.189* | 0.154* | 0.176* | 0.172* | 0.134* | 0.134* | 0.275* | 0.003* |

MOLECULAR ECOLOGY

**Table S3.** $F_{ST}$ among adult and seedling cohorts across six populations, and between high and low elevation population pairs in seedlings calculated using SPAGeDi. *: p-value < 0.01 from 10000 permutations.

| Population | $F_{ST}$ among cohorts | $F_{ST}$ high-low |
|---|---|---|
| **APE_H** | 0.007* | 0.018* |
| **APE_L** | 0.007* | |
| **BAV_H** | 0.002 | 0.010* |
| **BAV_L** | 0.000 | |
| **FAG_H** | 0.002* | 0.010* |
| **FAG_L** | -0.002* | |

MOLECULAR ECOLOGY

**Table S4.** Multi-class test statistic (ω, lower panel) and p-values (upper panel) for each pair of populations. Significant heterogeneity in the spatial autocorrelation in allele frequencies between populations are in bold (p-values < 0.01, following Banks & Peakall, 2012).

| | VEN_H | VEN_L | VES_H | VES_L | APE_H | APE_L | BAV_H | BAV_L | FAG_H | FAG_L | ISS_H | ISS_L | LUR_H | LUR_L | PYR_H | PYR_L |
|---|---|---|---|---|---|---|---|---|---|---|---|---|---|---|---|---|
| **VEN_H** | | 0.01 | 0.51 | 0.97 | 1.00 | 0.10 | 0.73 | 0.68 | 0.89 | 0.64 | 0.76 | 0.97 | 0.96 | 0.96 | 0.54 | 0.97 |
| **VEN_L** | 35.8 | | 0.90 | 0.82 | 1.00 | 0.08 | 0.61 | 0.49 | 0.77 | 0.92 | 1.00 | 1.00 | 0.99 | 0.96 | 0.50 | 0.72 |
| **VES_H** | 19.1 | 12.5 | | 0.42 | 0.21 | **0.00** | 0.67 | 0.29 | 0.47 | 0.86 | 0.22 | 0.56 | 0.14 | 0.76 | 0.04 | 0.70 |
| **VES_L** | 10.4 | 14.6 | 20.1 | | 0.38 | **0.00** | 0.77 | 0.47 | 0.72 | 0.57 | 0.74 | 0.93 | 0.05 | 0.90 | **0.00** | 0.14 |
| **APE_H** | 5.5 | 4.2 | 25.0 | 21.3 | | 0.03 | 0.89 | 0.96 | 1.00 | 0.99 | 0.33 | 0.60 | 0.99 | 0.62 | 0.93 | 0.98 |
| **APE_L** | 28.5 | 29.4 | **58.8** | **46.6** | 33.9 | | 0.06 | 0.17 | 0.01 | 0.16 | **0.00** | **0.00** | 0.06 | **0.00** | 0.02 | 0.09 |
| **BAV_H** | 15.7 | 17.3 | 16.6 | 15.1 | 12.7 | 30.7 | | 0.90 | 0.87 | 0.92 | 0.63 | 0.71 | 0.72 | 0.80 | 0.30 | 0.68 |
| **BAV_L** | 16.6 | 19.6 | 22.7 | 19.8 | 10.9 | 26.1 | 12.4 | | 0.53 | 0.98 | 0.30 | 0.46 | 0.71 | 0.55 | 0.71 | 0.86 |
| **FAG_H** | 13.1 | 15.2 | 19.4 | 15.9 | 6.7 | 36.9 | 12.9 | 18.6 | | 0.66 | 0.65 | 0.92 | 0.98 | 0.90 | 0.91 | 0.70 |
| **FAG_L** | 17.2 | 11.7 | 18.4 | 13.7 | 8.5 | 26.0 | 11.6 | 9.8 | 16.7 | | 0.74 | 0.82 | 0.93 | 0.72 | 0.53 | 0.88 |
| **ISS_H** | 15.4 | 6.3 | 11.4 | 24.9 | 22.2 | **49.0** | 17.3 | 22.5 | 17.1 | 15.8 | | 0.98 | 0.15 | 0.81 | 0.01 | 0.10 |
| **ISS_L** | 10.3 | 7.5 | 7.8 | 17.4 | 17.9 | **52.9** | 16.1 | 19.9 | 12.0 | 14.5 | 5.7 | | 0.34 | 1.00 | 0.08 | 0.32 |
| **LUR_H** | 10.3 | 7.8 | 31.5 | 26.2 | 8.3 | 30.5 | 15.8 | 16.2 | 10.1 | 11.8 | 26.2 | 21.9 | | 0.30 | 0.62 | 0.76 |
| **LUR_L** | 10.6 | 10.6 | 9.0 | 14.1 | 17.7 | **54.2** | 14.4 | 18.7 | 12.5 | 16.0 | 11.0 | 3.6 | 22.5 | | 0.02 | 0.31 |
| **PYR_H** | 18.6 | 19.2 | **41.9** | 31.9 | 11.8 | 36.0 | 22.7 | 16.4 | 12.1 | 18.7 | 38.4 | 29.2 | 17.8 | 33.5 | | 0.42 |
| **PYR_L** | 10.2 | 16.0 | 26.5 | 16.2 | 9.4 | 28.5 | 16.6 | 13.5 | 16.0 | 13.0 | 28.4 | 22.5 | 15.3 | 22.5 | 20.6 | |

MOLECULAR ECOLOGY

**Table S5.** Multi-class test statistic (ω, lower panel) and p-values (upper panel) between adults and saplings. Significant heterogeneity in the spatial autocorrelation in allele frequencies between populations are in bold (p-values < 0.01, following Banks & Peakall, 2012).

| | **APE_H** | **APE_H seedling** | **APE_L** | **APE_L seedling** |
|---|---|---|---|---|
| **APE_H** | | 0.64 | **0.00** | 0.91 |
| **APE_H seedling** | 17.2 | | **0.00** | 0.44 |
| **APE_L** | **40.7** | **44.6** | | 0.06 |
| **APE_L seedling** | 12.0 | 20.3 | 31.1 | |

| | **BAV_H seedling** | **BAV_H** | **BAV_L seedling** | **BAV_L** |
|---|---|---|---|---|
| **BAV_H seedling** | | 0.02 | **0.00** | **0.00** |
| **BAV_H** | 34.2 | | 0.48 | 0.56 |
| **BAV_L seedling** | **47.1** | 19.3 | | 0.01 |
| **BAV_L** | **61.0** | 18.5 | 37.8 | |

MOLECULAR ECOLOGY

**Table S6.** Sensitivity of *Sp* to the spatial configuration of the sampling the two VEN populations, and with three resampling methods. See Materials and Methods for more details. Sig *Sp* is the percentage of significant *Sp* values. Mean and SD of *Sp* values after the 20 resampling, and result of One-Sample t-test between the *Sp* values of the resamplings and the original *Sp* values (0.006 and 0.003 for VEN_H and VEN_L, respectively).

| | Resampling method | Sig *Sp* (%) | Mean *Sp* ± SD | One-Sample t-test | |
|---|---|---|---|---|---|
| | | | | t | p-value |
| **VEN_H** | random | 60 | 0.006 ± 0.003 | 0.096 | 0.924 |
| | reduced area | 85 | 0.006 ± 0.002 | 0.252 | 0.804 |
| | transect | 95 | 0.006 ± 0.002 | -0.308 | 0.761 |
| **VEN_L** | random | 10 | 0.002 ± 0.002 | -0.479 | 0.637 |
| | reduced area | 80 | 0.007 ± 0.004 | 6.132 | 0.000 |
| | transect | 80 | 0.005 ± 0.002 | 5.664 | 0.000 |

MOLECULAR ECOLOGY

**Table S7.** Result of the structural equation modelling (SEM) analyses of *Sp* and demography with the best model.

Initial and best model:
**'*Sp* ~ High/Low + LD1.DAPC + LD2.DAPC'**

| | |
|---|---|
| ***Model Test User Model*** | |
| Test statistic | 0.000 |
| Degrees of freedom | 0 |
| ***Model Test Baseline Model*** | |
| Test statistic | 10.08 |
| Degrees of freedom | 3 |
| P-value | 0.018 |
| ***User Model vs Baseline Model*** | |
| Comparative Fit Index (CFI) | 1.000 |
| Tucker-Lewis Index (TLI) | 1.000 |
| ***Loglikelihood and Information Criteria*** | |
| Loglikelihood user model (H0) | 73.00 |
| Loglikelihood unrestricted model (H1) | 73.00 |
| Akaike (AIC) | -138.00 |
| Bayesian (BIC) | -134.91 |
| Sample-size adjusted Bayesian (BIC) | -147.15 |
| ***Root Mean Square Residuals*** | |
| RMSEA | 0.000 |
| 90 Percent confidence interval lower | 0.000 |
| 90 Percent confidence interval upper | 0.000 |
| P-value RMSEA<=0.05 | NA |
| ***Standardized Mean Square Residuals*** | |
| SRMR | 0.000 |
| ***R-Square:*** | |
| ***Sp*** | **0.468** |

| | | **Estimate** | **Std.Err** | **z-value** | **P(>\|z\|)** | **Std.lv** | **Std.all** |
|---|---|---|---|---|---|---|---|
| ***Regressions:*** | | | | | | | |
| *Sp~* | | | | | | | |
| | High/Low | 0.002 | 0.001 | 2.794 | 0.005 | 0.005 | 0.510 |
| | LD1.DAPC | -0.000 | 0.000 | -2.289 | 0.022 | -0.000 | -0.418 |
| | LD2.DAPC | -0.000 | 0.000 | -0.867 | 0.386 | -0.000 | -0.158 |
| ***Variances:*** | | | | | | | |
| | *.Sp* | 0.000 | 0.000 | 2.828 | 0.005 | 0.000 | 0.532 |

MOLECULAR ECOLOGY

**Table S8.** Result of the structural equation modelling (SEM) analyses of *Sp* and environmental variables with the best model.

| | |
|---|---|
| Initial model: | |
| '*Sp* ~ Temperature + Isothermality + Precipitation + PC1.spcomp + PC2.spcomp + PC3.spcomp + AWC + Elevation | |
| PC1.spcomp ~ Temperature + Isothermality + Precipitation + AWC + Elevation | |
| PC2.spcomp ~ Temperature + Isothermality + Precipitation + AWC + Elevation | |
| PC3.spcomp ~ Temperature + Isothermality + Precipitation + AWC + Elevation' | |
| Best model: | |
| **'*Sp* ~ Isothermality + PC2.spcomp + AWC + Elevation** | |
| **PC2.spcomp ~ Temperature + Precipitation + AWC + Elevation'** | |
| ***Model Test User Model*** | |
| Test statistic | 0.880 |
| Degrees of freedom | 3 |
| P-value (Chi-square) | 0.830 |
| ***Model Test Baseline Model*** | |
| Test statistic | 40.33 |
| Degrees of freedom | 11 |
| P-value | 0.000 |
| ***User Model vs Baseline Model*** | |
| Comparative Fit Index (CFI) | 1.000 |
| Tucker-Lewis Index (TLI) | 1.265 |
| ***Loglikelihood and Information Criteria*** | |
| Loglikelihood user model (H0) | 58.95 |
| Loglikelihood unrestricted model (H1) | 59.39 |
| Akaike (AIC) | -97.90 |
| Bayesian (BIC) | -90.17 |
| Sample-size adjusted Bayesian (BIC) | -120.77 |
| ***Root Mean Square Residuals*** | |
| RMSEA | 0.000 |
| 90 Percent confidence interval lower | 0.000 |
| 90 Percent confidence interval upper | 0.246 |
| P-value RMSEA<=0.05 | 0.839 |
| ***Standardized Mean Square Residuals*** | |
| SRMR | 0.019 |
| ***R-Square:*** | |
| ***Sp*** | **0.582** |
| **PC2.spcomp** | **0.804** |

| | | Estimate | Std.Err | z-value | P(>\|z\|) | Std.lv | Std.all |
|---|---|---|---|---|---|---|---|
| ***Regressions:*** | | | | | | | |
| *Sp~* | | | | | | | |
| | Elevation | 0.002 | 0.001 | 3.258 | 0.001 | 0.002 | 0.646 |
| | Isothermality | 0.002 | 0.001 | 3.154 | 0.002 | 0.002 | 0.542 |
| | AWC | -0.001 | 0.001 | -1.999 | 0.046 | -0.001 | -0.382 |
| | PC2.spcomp | 0.001 | 0.000 | 2.370 | 0.018 | 0.001 | 0.461 |
| PC2.spcomp~ | | | | | | | |
| | Elevation | -1.533 | 0.213 | -7.185 | 0.000 | -1.533 | -1.018 |
| | Temperature | -1.071 | 0.197 | -5.436 | 0.000 | -1.071 | -0.711 |
| | Precipitation | 0.527 | 0.178 | 2.956 | 0.003 | 0.527 | 0.350 |
| | AWC | 0.970 | 0.188 | 5.173 | 0.000 | 0.970 | 0.644 |
| ***Variances:*** | | | | | | | |
| | .*Sp* | 0.000 | 0.000 | 2.828 | 0.005 | 0.000 | 0.418 |
| | .PC2.spcomp | 0.418 | 0.148 | 2.828 | 0.005 | 0.418 | 0.196 |

MOLECULAR ECOLOGY

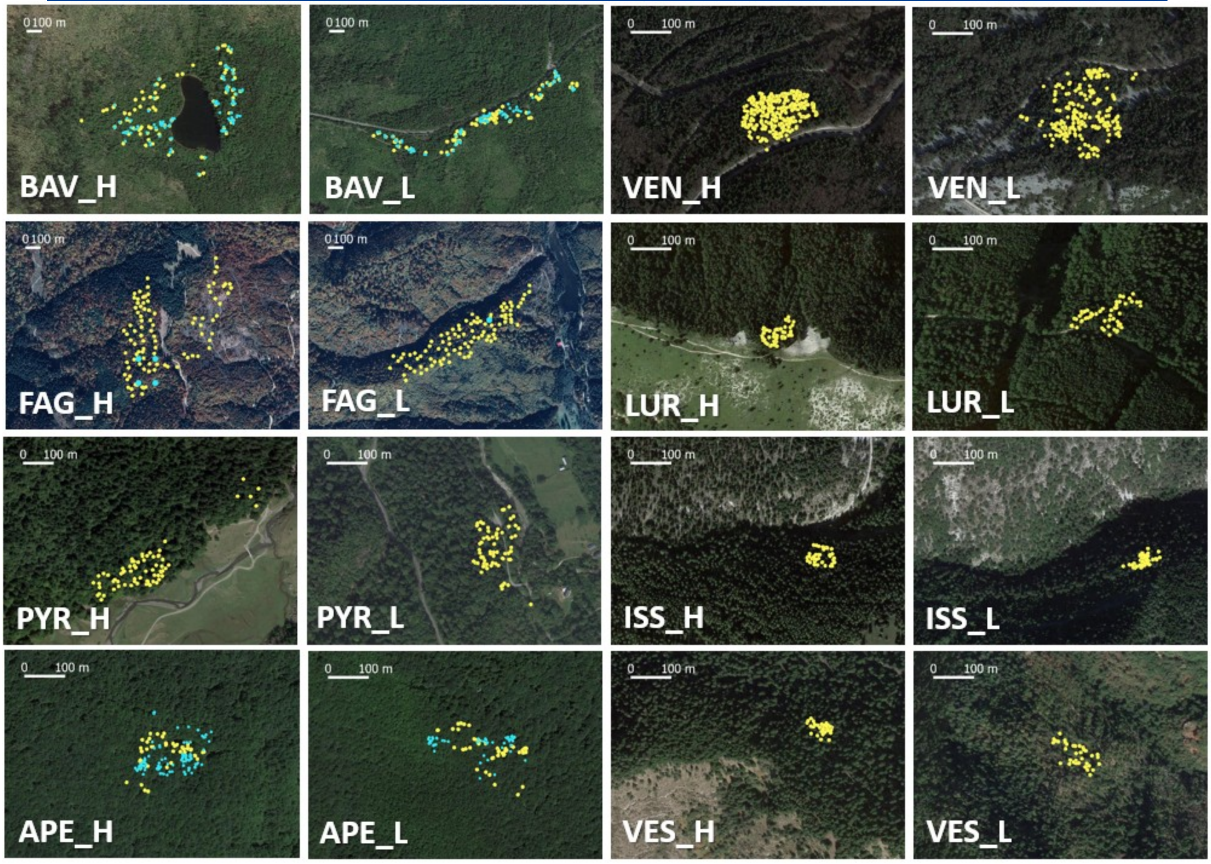


**Figure S1.** Location of each sampled tree in each population separately. Adult trees are marked with yellow dots and the seedlings by blue dots.

MOLECULAR ECOLOGY

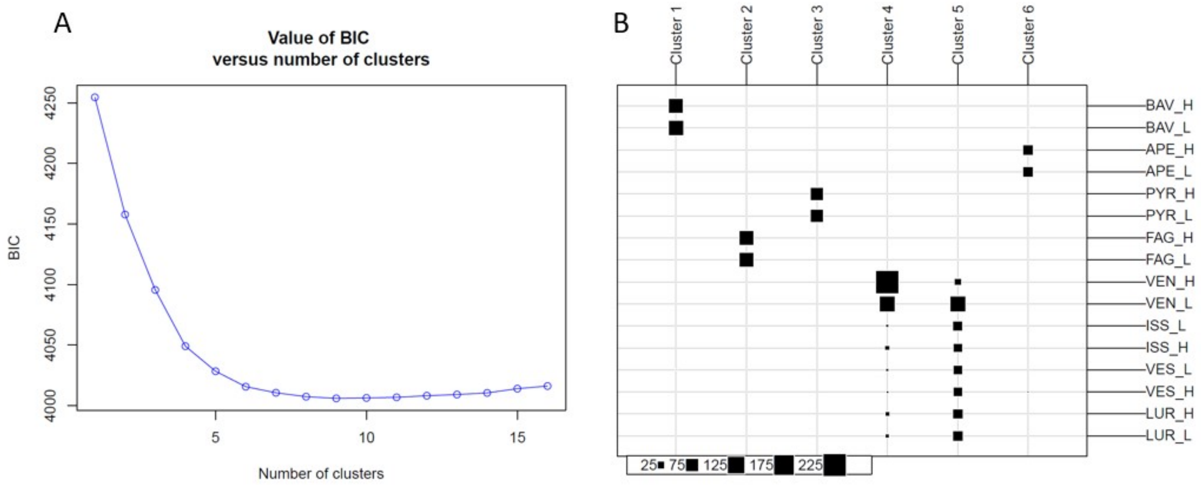


**Figure S2.** Population structure revealed by the discriminant analysis of principal components (DAPC): (A) The optimal number of clusters (K) as determined by ‘k-means’, Bayesian Inference Criterion (BIC) values was represented versus against number of clusters and (B) number of clusters was represented against populations.

MOLECULAR ECOLOGY

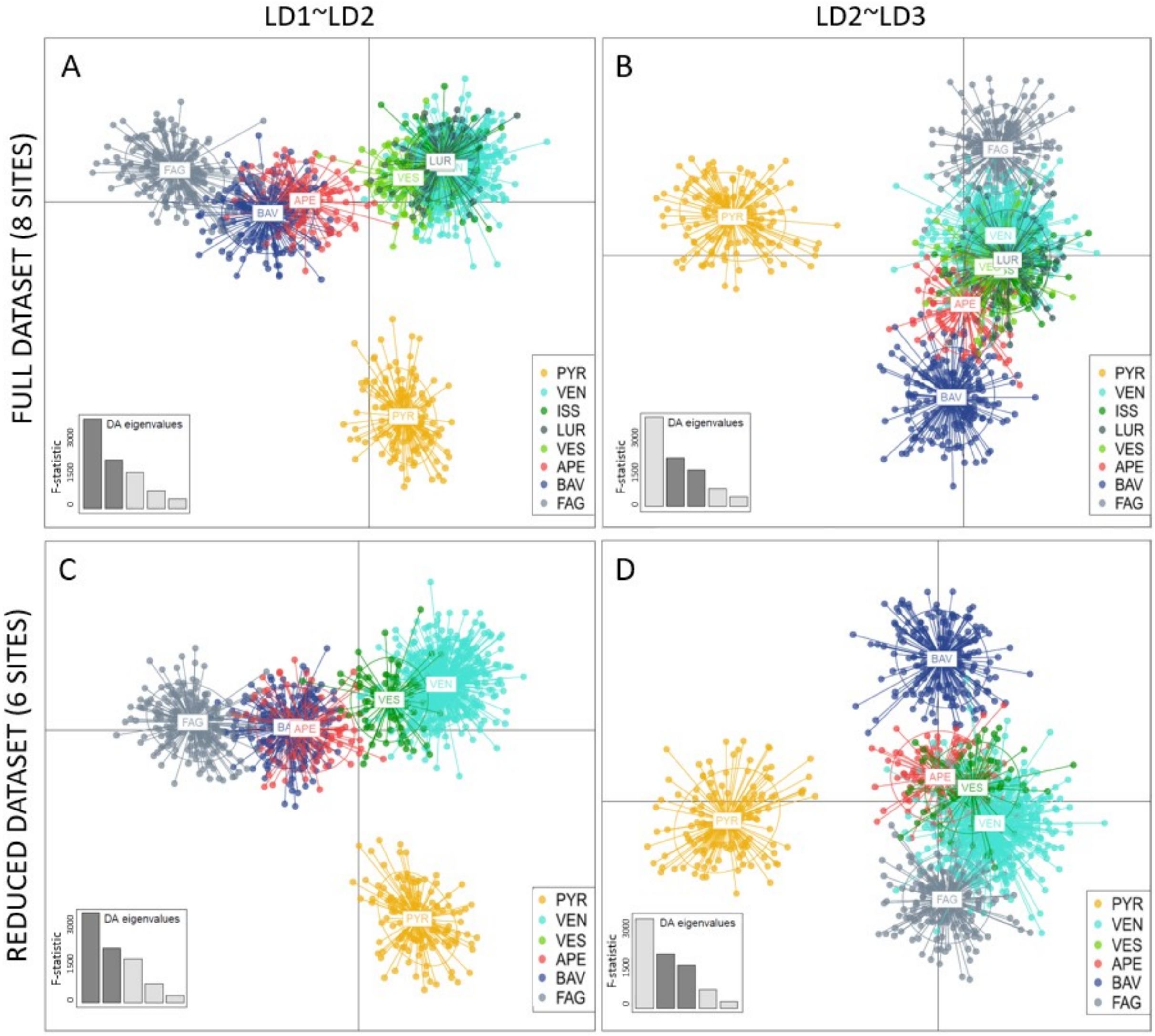


**Figure S3.** Population structure revealed by the discriminant analysis of principal components (DAPC): Scatterplot based on the DAPC output for six assigned genetic clusters, sites indicated by different colours. Dots represent different individuals. (A) DAPC based on all studied population, representing the first two Linear Discriminants (LD) and (B) representing the second and third LD. (C) DAPC based on six sites (only VEN and VES was included from the Western Alps), representing the first two LD and (C) representing the second and third LD.

MOLECULAR ECOLOGY

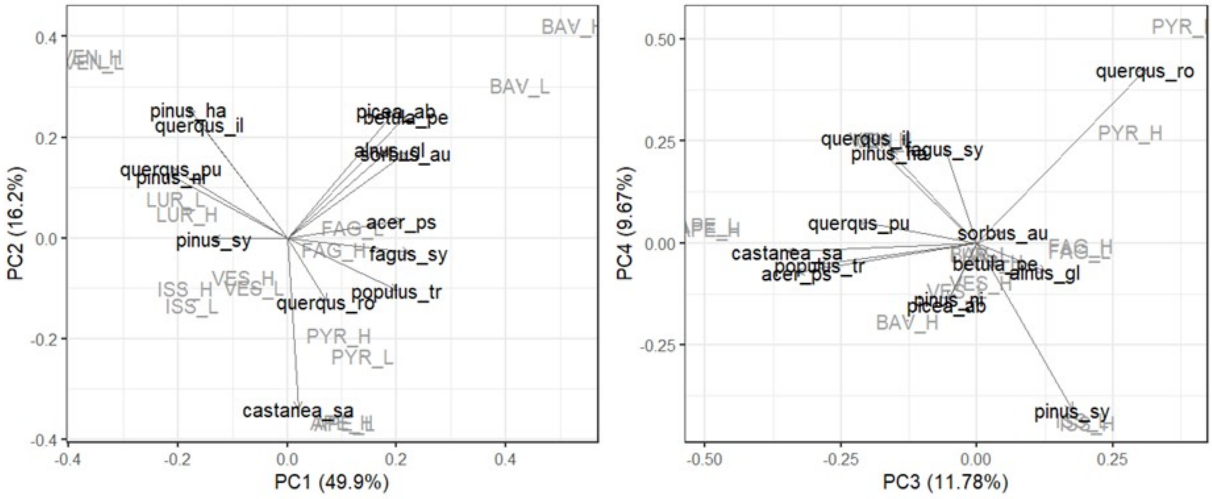


**Figure S4.** Situation of the 16 population locations along the first four Principal Components of species composition using 14 tree species (*Acer pseudoplatanus*, *Alnus glutinosa*, *Betula pendula*, *Castanea sativa*, *Fagus sylvatica*, *Picea abies*, *Pinus halapensis*, *Pinus nigra*, *Pinus sylvestris*, *Populus tremula*, *Querqus ilex*, *Querqus pubescens*, *Querqus robur*, *Sorbus aucuparia*).

MOLECULAR ECOLOGY

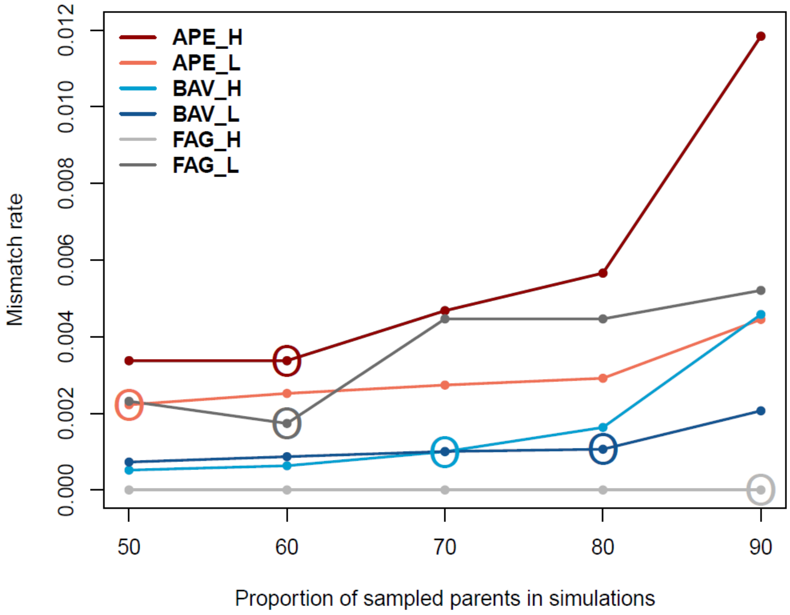


**Figure S5.** Representation of mismatch rates with different proportions of sampled parents (50. 60. 70. 80 and 90 %) used in the simulations for the parentage analyses. The circled dots indicate the proportions of sampled parents selected for the final analyses.
